# Cryo-EM image processing of amyloid filaments in RELION-5.1

**DOI:** 10.64898/2026.03.17.712386

**Authors:** Sofia Lövestam, Jenny Shi, David Li, Kiarash Jamali, Sjors H.W. Scheres

## Abstract

We present new tools for the structure determination of amyloid filaments from electron cryo-microscopy (cryo-EM) images. We introduce a new algorithm for automated filament picking, based on their characteristic 4.75 Å repeat signal; we implement the new auto-picker in a fully automated procedure for on-the-fly pre-processing of cryo-EM data sets of amyloid filaments; we present a graphical tool to select filament types based on bi-hierarchical clustering of filaments and 2D class assignments; and we introduce a denoising neural network for Blush regularisation that is re-trained on amyloid reconstructions. The implementation of these tools in release 5.1 of our open-software package RELION ensures broad applicability. We demonstrate their usefulness on two experimental data sets, including a previously described data set on recombinant human islet amyloid protein (hIAPP) with the S20G mutation for which we identify two new filament types.

## Introduction

More than a decade after the electron cryo-microscopy (cryo-EM) resolution revolution ^1^, electron microscopy has become the dominant technique for atomic structure determination of biological macromolecular complexes ^2^. Fuelled by improved signal-to-noise performance of direct electron detectors ^3^, the correction of beam-induced motions by the processing of movies ^4,5^ and enhanced particle alignment via regularised likelihood optimisation ^6^, cryo-EM has enabled atomic structure determination for a wide range of samples that were hitherto intractable for structural biology, including amyloid filaments.

Amyloid filaments are helical protein assemblies, where multiple copies of a given protein (or of various proteins) stack on top of each other, forming long β-sheets along the direction of their helical axis. Many amyloid filaments are associated with disease ^7^, most notably in neurodegeneration ^8^, whereas others serve physiological roles ^9^, such as in bacterial biofilm formation ^10^.

Prior to 2017, atomic structure determination of amyloid filaments was limited to electron or X-ray crystallography or solid-state nuclear magnetic resonance techniques ^11^. Due to the absence of low-resolution structural features along their helical axis, structure determination of amyloids by cryo-EM was hampered by alignment issues. Consequently, the resolution of cryo-EM reconstructions of amyloids precluded *de novo* generation of atomic models ^12,13^. This changed with the implementation of helical reconstruction in the RELION software ^14^, which combined the iterative helical real-space refinement (IHRSR) approach ^15^ with regularised likelihood optimisation ^6^. This combination enabled the reconstruction of tau amyloid filaments from the brain of an individual with Alzheimer’s disease to resolutions sufficient for *de novo* atomic modelling ^16^. Since then, many amyloid structures have been determined by cryo-EM, as documented in the Amyloid Atlas ^17^, which currently refers to more than 500 amyloid structures in the Electron Microscopy Data Bank (EMDB) ^18^.

It is therefore unsurprising that, in recent years, numerous methodological advances have been developed for the cryo-EM analysis of amyloid filaments. These include approaches for automated filament picking in micrographs ^19–22^; for the separation of filament types in structurally heterogeneous data sets ^23–25^; for generating initial reference models to bootstrap three-dimensional refinement ^26^; and for the application of additional restraints on segment orientations within individual filaments ^27^. Despite these advances, cryo-EM structure determination of amyloid filaments remains challenging. The absence of low-resolution features along the helical axis, combined with the tendency of helical reconstruction to become trapped in incorrect local minima ^28^, makes amyloid reconstruction more prone to erroneous solutions than the reconstruction of globular complexes. Moreover, because a given protein can adopt multiple distinct amyloid conformations, cryo-EM data sets frequently contain mixtures of different filament types, which need to be separated to allow reliable reconstruction ^20^.

Here, we describe recent developments for the processing of amyloids in RELION. We introduce a new approach for the automated picking of amyloid filaments in cryo-EM micrographs based on their characteristic 4.75 Å signal in the Fourier domain; an approach that builds on the Clustering of HElical Polymers (CHEP) algorithm ^24,25^ to identify and separate filament types according to their contributions to 2D class averages; a denoising neural network trained specifically to regularise amyloid reconstructions; and an automated pipeline for micrograph pre-processing up to the stage of 2D class averaging. We demonstrate the utility of these developments using two previously published data sets: one comprising filaments formed from recombinant tau with twelve phosphomimetic mutations (PAD12) adopting the paired helical filament fold, and another data set from a time-resolved study on the *in vitro* assembly of human islet amyloid polypeptide (hIAPP) ^29^.

## Approach

### An amyloid-specific auto-picker

Because successive rungs along the β-sheets of amyloids repeat every ∼4.75 Å, cryo-EM images of amyloids have a characteristic signal in their Fourier transform. We developed an algorithm that detects, quantifies and determines the orientation of this signal in micrographs. To minimise computational costs, micrographs are downscaled to a pixel size of 2.1 Å, and for every fifth downscaled pixel (in both X and in Y), we extract 1D arrays of neighbouring pixels in 36 uniformly sampled directions (i.e. sampled every 5°). The length of the 1D arrays is user-defined (by default 1D arrays are 250 Å long). For each array, we compute 1D Fourier transforms and calculate the accumulated power within the resolution range of 4.65 and 4.85 Å. To boost the signal-to-noise ratio, the accumulated power values from neighbouring, parallel 1D arrays within 50 Å are averaged.

For each sampled pixel, we calculate a Z-score by dividing the maximum accumulated power across the 36 directions by the standard deviation of the remaining 33 directions, excluding the direction with the maximum power and its two neighbouring directions. The image with the Z-scores for all sampled, downscaled pixels, which we call the figure-of-merit (FOM) image, is written to disk, alongside a corresponding PSI image that contains the in-plane rotation angles at which the accumulated power in the 4.65-4.85 Å range is maximal. Because ice crystals in the micrographs can lead to spurious Z-scores in this resolution range, we also calculate Z-scores for the accumulated power between 4.2-4.4 Å. Any pixels for which the latter Z-scores are higher than 0.7 are considered likely to arise from ice contamination and are therefore set to zero in the FOM image.

The combined FOM and PSI images are fed into a neural network with a modified U-net architecture ^30^. The modifications include using instance normalisation ^31^ and a Gaussian Error Linear Unit for the activation function ^32^. Instance normalisation is better than batch normalisation for cryo-EM image processing due to the expected, large differences of pixel values across batches (Jamali et al., 2024). Batch normalisation averages these differences across batches, causing meaningful signal to be lost.

For training this neural network, roughly 2,000 micrographs containing tau, α-synuclein, amyloid-β, TDP43 and TMEM106B filaments were gathered, FOM and PSI images were calculated, and filaments manually identified. During training, we provide the network with the FOM and PSI image pairs, from which it must output a predicted probability that each pixel contains a filament. We then calculate the binary cross entropy loss between the predicted and target images.

During inference, FOM and PSI images of micrographs are similarly calculated and provided to the trained neural network. The predicted segmentation mask from the network is binarized at a user-specified threshold, and the binary image is skeletonized using the scikit-image Python library ^33,34^. The resulting skeletons are divided into individual paths using the Skeleton function from the skan python library ^35^. Finally, branches, paths that are within a user-defined minimum distance from each other and paths shorter than a user-defined minimum length are removed. The resulting paths are written in a new RELION STAR file format that represents curved filaments as sequences of linear segments of arbitrary length. The “Particle extraction” jobtype in RELION reads these STAR files to extract individual filament segment images at a user-defined distance from each other along these lines.

### Automated pre-processing

The new auto-picker has been incorporated in an automated pre-processing procedure that is specific for micrographs with amyloid filaments. Based on the Schemes introduced in RELION-4.0 ^36^, a first Scheme called *amyprep* iteratively executes cycles of “Import”, “Motion Correction”, “CTF estimation”, “Auto-picking” and “Subset selection” jobs, in a project directory where new micrograph movies are copied as they are being collected. (The Scheme can be set up and executed using the command “relion_schemegui amyprep”). The Auto-picking job in the amyprep Scheme computes the FOM/PSI maps but does not call the neural network for filament tracing. Thereby, the entire amyprep Scheme may be executed on a computer that has been optimised for CPU execution. The subset selection job selects all micrographs for which the skewness of the observed FOM values is larger than 1, which effectively removes micrographs without amyloid filaments from the data set.

The second Scheme, called *amyproc*, runs in parallel with amyprep and may be executed on a computer that is optimised for GPU calculations. (One could also run both Schemes on the same computer.) This Scheme calls a second Auto-picking job that reads in the FOM/PSI images from the Auto-Picking job of the amyprep Scheme and uses the neural network to trace filaments in the micrographs. This job is then followed by “Particle Extraction”, “Subset selection” and “2D classification” jobs, where the latter are run in batches of a user-defined number of helical segments. By setting the batch_size parameter to a value larger than the number of particles, all segments are combined into a single 2D classification.

### A graphical tool to identify and select filament types

Our filament type selection tool is conceptually related to the CHEP algorithm ^24,25^, which exploits the observation that individual particle segments from a given filament in most cases belong to the same filament type, i.e. transitions between filament types within a single filament are rare. In 2D classification of particle images this information is not used, and individual particle image segments are separated independently into multiple classes. Because segments from different filament types will distribute differently among the 2D classes, a (2D class ID × filament ID) particle counts matrix has an inherent modular structure. Segments along a given filament are a sample from the distribution of their relevant filament type. Whereas the CHEP algorithm uses k-means clustering of the particle counts matrix, followed by PCA analysis, we use a bi-hierarchical clustering method, clustermap from the seaborn Python library ^37^, with cosine distance as the similarity metric, and visualise the clustered particle counts matrix using fcluster from the SciPy Python library ^38^. In this matrix, different filament types represent blocks when filament IDs and 2D class IDs are grouped correctly (see Figures 1-2). The hierarchical clustering of 2D class averages also reorders the 2D class averages so that averages found in similar sets of filaments are placed closer together, providing a direct visual grouping of classes, analogous to the sorting one would do for different filament types in conventional manual selection of 2D class average images. This tool thereby provides a convenient means to visually assess the structural heterogeneity of the data, and to select subsets of the data corresponding to different filament types based on selecting neighbouring columns in the clustered particle counts matrix. Repeated application of the tool in tandem with 2D classification can further separate filament types, including rare filament types present initially in only a few 2D classes as demonstrated in our previous work using a development version of the tool prior to RELION integration ^39^.

**Figure 1:**
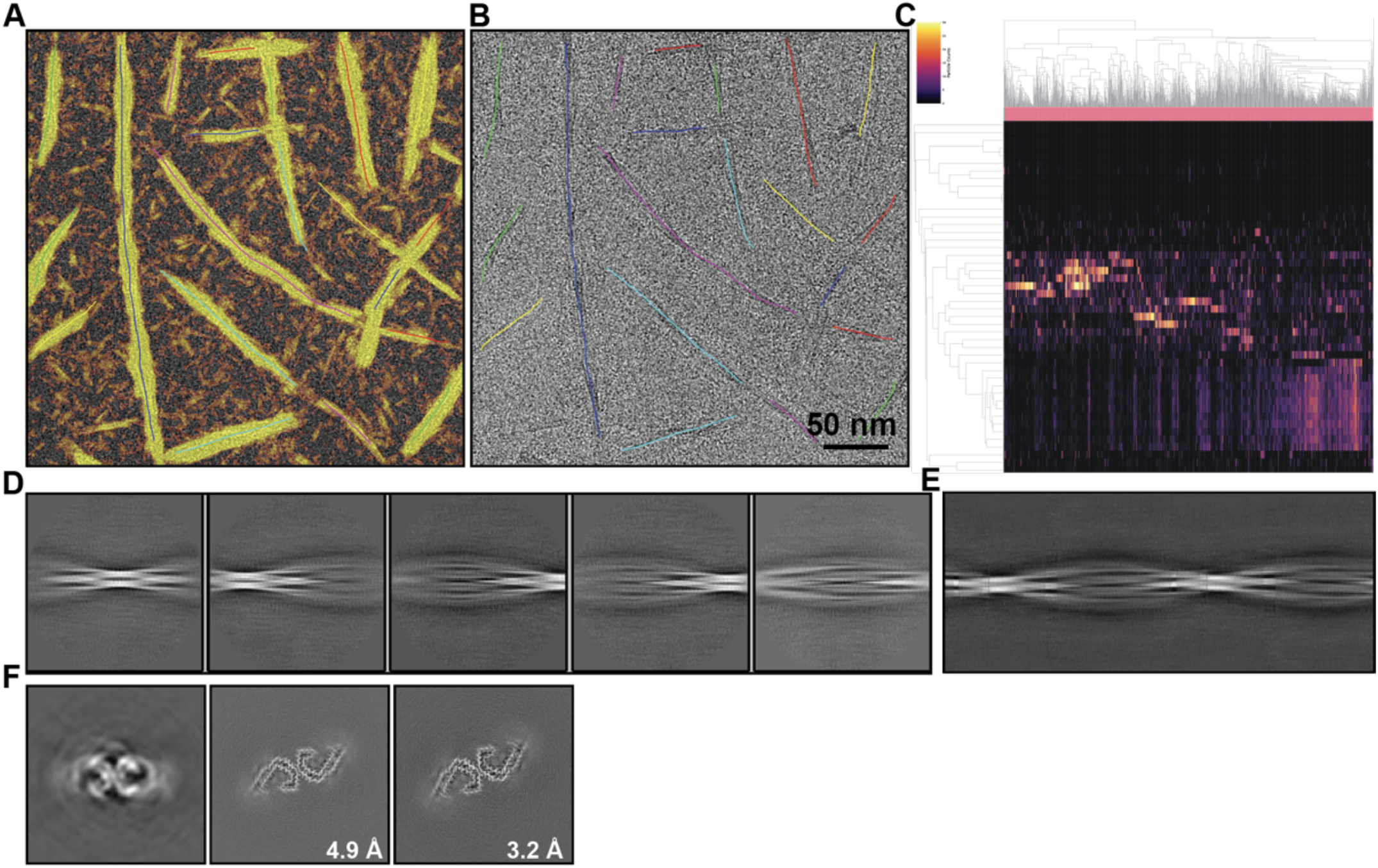
A recombinant PAD12 tau data set. **(A)** Micrograph coloured by the FOM signal (from red to yellow for increasing FOM values above 0.5). **(B)** Auto-picked filaments. **(C)** Bi-hierarchical clustering after 2D classification. Filaments are ordered horizontally; 2D classes are ordered vertically. **(D)** 2D class averages selected for initial model generation. **(E)** Image of a full cross-over obtained by relion_helix_inimodel2d. **(F)** Slices through the initial model (left), the map after the first refinement with amyloid-specific Blush regularisation, and the final map (right).

**Figure 2:**
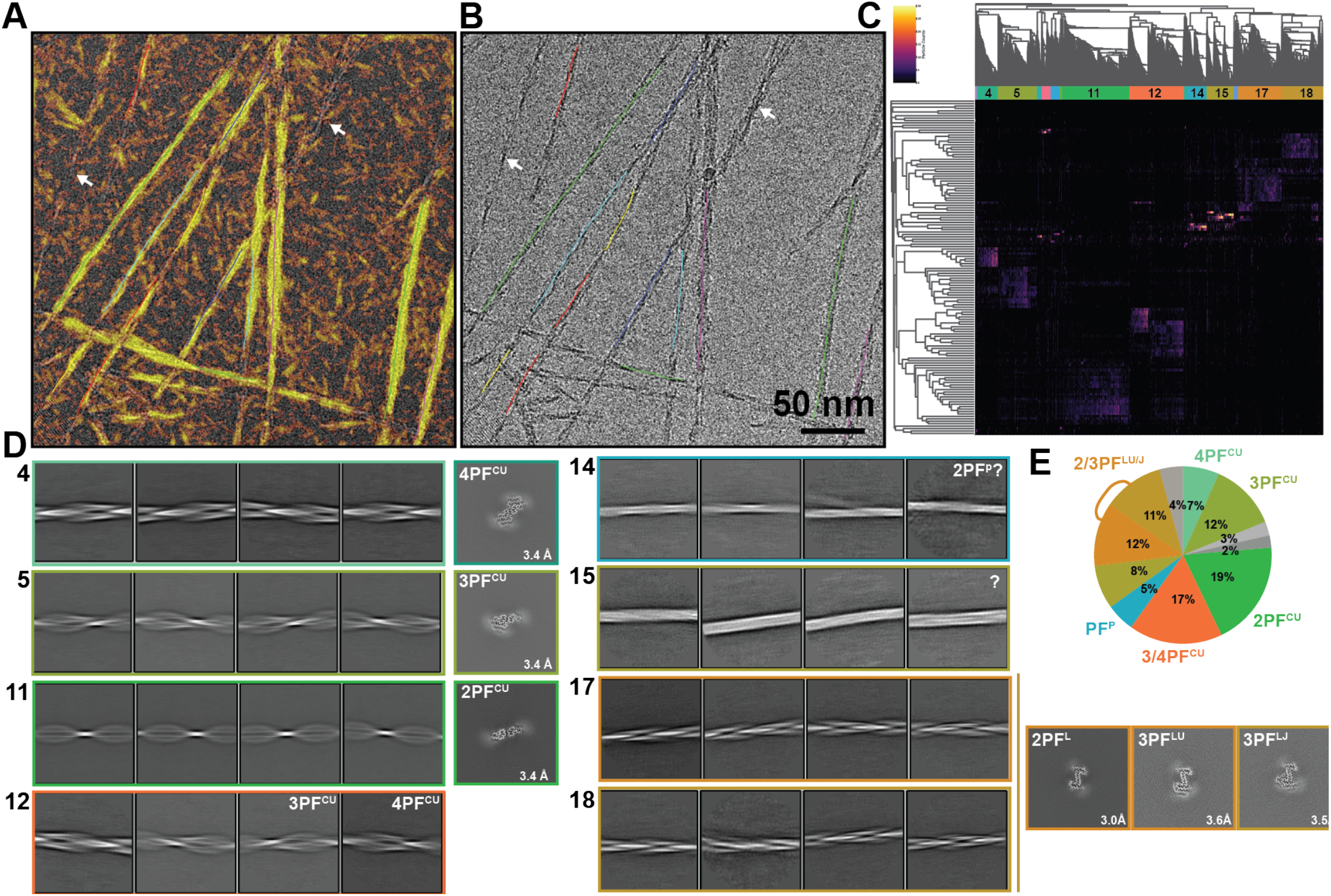
A recombinant hIAPP S20G data set. **(A)** Micrograph coloured by the FOM signal (from red to yellow for increasing FOM values above 0.5). White arrows indicate filaments without strong FOM signals. **(B)** Auto-picked filaments. **(C)** Bi-hierarchical clustering after 2D classification. Filaments are ordered horizontally; 2D classes are ordered vertically. **(D)** 2D class averages selected for each of the eight clusters identified in the bi-hierarchical clustering (left), plus slices through their corresponding 3D reconstructions, if successful (right). **(E)** Percentages of the particles in each of the clusters from the bi-hierarchical clustering. The eight selected filaments are coloured with the same colours as in panels C and D; clusters with images unsuitable for further processing are shown in grey.

### An amyloid-specific network for Blush regularisation

We previously described Blush regularisation as a powerful way to incorporate prior knowledge about cryo-EM maps in the reconstruction process ^40^. Blush regularisation uses a denoising neural network that was trained on 422 pairs of reconstructions from cryo-EM half-sets from the Electron Microscopy Data Bank (EMDB) ^18^. Although this network did improve the reconstruction of the first intermediate amyloid in the assembly of tau (297-391) into paired helical filaments ^39^, it incorporates no specific knowledge about amyloid reconstructions as it was trained on a general set of structures from the EMDB. We have since experienced that the general Blush network often fails to improve reconstructions of amyloids for difficult data sets.

Therefore, we optimised the weights of the general Blush network for its specific use on reconstructions of amyloid filaments. We downloaded 505 PDB entries that were listed on the Amyloid Atlas ^17^ on 13 Feb 2025. Of these, we identified 361 entries that had reconstructions from half-sets in the EMDB. Manual inspection of these entries was used to discard suboptimal reconstructions and to correct errors in the deposited metadata values for the helical twist and rise, resulting in a final training set of 318 entries (Supplementary Table 1).

We used the atomic models that were associated with the EMDB entries, together with the reported helical twist and rise values, to generate tight binary masks around those parts of the reconstructions that were suitable for atomic modelling. The binary masks were extended with a soft raised cosine edge with a width of 7.5 Å using relion_image_handler.

The resulting combinations of half-set reconstructions and soft-edged masks were then used to re-train the previously reported Blush model ^40^, with the following modifications. We restricted augmentation of the rotations to 90-degree rotations around the helical axis only, so that the network could learn about the amyloid-specific signal that repeats every ∼4.75 Å along the helical axis, and the fact that amyloid reconstructions extend all the way until the edge of the box in the Z-direction. We also skipped the use of anisotropy in the augmentation of the map filters, because amyloid reconstructions are typically calculated from uniformly sampled orientations around the helical axis. Like we did for the training of the general Blush network, we used the same lower resolution cut-offs for low-pass filtering half-set map pairs to minimise the risk of hallucinating high-resolution features by the denoiser.

Starting from the weights of the published Blush model, we re-trained an amyloid-specific denoiser network for 170,000 steps with a learning rate of 2e-5; a drop-out rate of 50%; and a batch size of 16. Training on four NVIDIA A100 GPUs took three days.

## Results

### A straightforward data set on recombinant PAD12 tau

We first applied the approaches described above on a data set of 651 micrograph movies of recombinant PAD12 tau filaments ^41^. Running amyprep and amyproc on a machine with a 32-core AMD Ryzen Threadripper PRO 7975WX CPU and 4 NVIDIA GeForce RTX4090 GPUs took approximately 3 hours to completion of the 2D class averaging of all 476,817 auto-picked helical segments. Micrographs showed strong FOM signals (Figure 1A), in which most filaments seemed to be picked correctly by visual inspection (Figure 1B). All 2D class averages resembled projections of tau paired helical filaments with the Alzheimer fold, and bi-hierachical clustering identified all filaments as belonging to the same type (Figure 1C). The 2D class averages shown in Figure 1D were used to calculate a *de novo* three-dimensional (3D) initial model using the relion_helix_inimodel2d program (Figure 1E) ^26^. 3D auto-refinement of this model using amyloid-specific Blush regularisation led to a preliminary map to 4.9 Å resolution. Subsequent particle polishing, CTF refinement and 3D auto-refinements with amyloid-specific Blush regularisation led to a final map to 3.2 Å resolution.

### A more challenging data set on recombinant IAPP

We next applied these procedures to a more challenging data set. This data set is publicly available from the EMPIAR database ^42^ as entry 11715 and consists of 925 micrographs that were recorded as part of a time-resolved cryo-EM study on the assembly of recombinant human islet amyloid polypeptide (hIAPP) ^29^. hIAPP is a secreted hormone that assembles into amyloid filaments in the pancreas of more than 90% of individuals with type II diabetes. The mutation of serine 20 into glycine (S20G) is associated with early-onset familial type II diabetes. The data set was recorded during the growth phase of the *in vitro* assembly of recombinant S20G hIAPP, i.e. six weeks after initiation of the reaction. Four distinct filament structures were reported at resolutions ranging from 3.1-3.4 Å.

Again, we ran the amyprep and amyproc schemes to preprocess micrographs, auto-pick filaments and perform 2D class averaging without user intervention. Using the same Threadripper machine, 2D class averages were obtained for all 531,977 auto-picked segments within 30 hours. In contrast to the tau data set described above, not all hIAPP filaments showed strong FOM signals (e.g. see white arrows in Figure 2A,B) and therefore not all filaments were picked. Bi-hierarchical clustering of the filaments and 2D class averages suggested the presence of at least eight distinct clusters of filaments (Figure 2C).

Separate 2D classifications were performed for each of the eight identified clusters (Figure 2D), and the resulting 2D class averages were used to calculate 3D initial models for subsequent refinements using the relion_helix_inimodel2d program ^26^.

Subsequent high-resolution 3D auto-refinements with amyloid-specific Blush regularisation, Bayesian polishing and CTF refinement of clusters 4, 5 and 11 yielded reconstructions to resolutions of 3.4 Å of hIAPP filaments with 4, 3 or 2 protofilaments with a C-shaped conformation, respectively (4PF^CU^, 3PF^CU^, 2PF^CU^). The same structures, at nearly identical resolutions, were also described in ^29^.

Clusters 14 and 15 showed relatively poor 2D class averages that did not yield suitable 3D initial models for further refinement. The 2D class averages in cluster 14 resemble those of filaments with 2 protofilaments in the P-shaped conformation, which were described in ^29^ to be present 3 weeks after starting the assembly reaction.

The 2D class averages of clusters 17 and 18 looked similar and filaments from these two clusters were merged. Subsequent 3D classification yielded three different structures. Bayesian polishing, CTF refinement and 3D auto-refinements with amyloid-specific Blush yielded maps to resolutions of 3.0-3.6 Å (Figure 2D; Figure 3). One of these structures was described in ^29^ and consists of two protofilaments in an L-shaped conformation (2PF^L^). The other two structures have not been described previously. They both comprise three protofilaments: two protofilaments adopt the L-shaped conformation, the third one adopts either a U or a J-shaped conformation (3PF^LU^ and 3PF^LJ^, respectively) (Figure 3; Supplementary Table 2).

**Figure 3:**
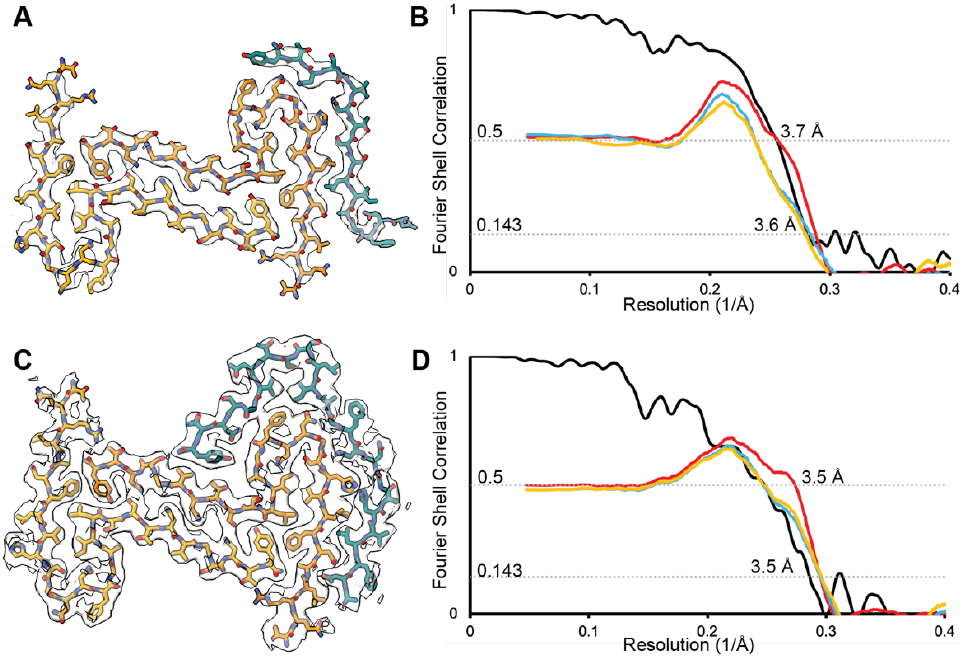
Structure of 3PF^LU^ and 3PF^LJ^. **(A)** Refined atomic model fitted into the cryo-EM density map of 3PF^LU^ (transparent grey). **(B)** Fourier Shell correlation curves between two independently refined half maps (black), the atomic model fitted in half-map 1 and half-map 1 (blue), between the atomic model fitted in half-map 1 and half-map 2 (yellow), and between the atomic model fitted in the sum of the two half-maps and the sum of the two half-maps (red) for 3PF^LU^. **(C)** as in (A) but for 3PF^LJ^. **(D)** as in (B) but for 3PF^LJ^.

### Filaments without FOM signals are of suboptimal quality

The observation that not all filaments of the IAPP data set had strong 4.75 Å FOM signals prompted us to investigate whether this represented a failure of the algorithm to identify what appeared to be good filaments by visual inspection of the micrographs. We manually traced filaments in the IAPP data set for which we did not observe FOM signals above 0.3. 2D classification of the manually traced filaments revealed they belonged primarily to the 2PF^L^ filament types. Comparison of 2D class averages of 33,030 segments from manually traced filaments without strong FOM signals with 33,252 2PF^L^ segments from auto-picking did not reveal obvious differences (Figure 4A,E). However, when we performed 3D auto-refinements starting from the same initial model for the two sets of segments, only those with strong FOM signals yielded a 3D reconstruction in which the layers of the amyloids were resolved (Figure 4B-D,I). In agreement with the absence of a strong 4.75 Å signal in the micrographs, the manually traced filaments led to a reconstruction with a resolution of 4.8 Å, which failed to separate the amyloid layers (Figure 4E-I).

**Figure 4:**
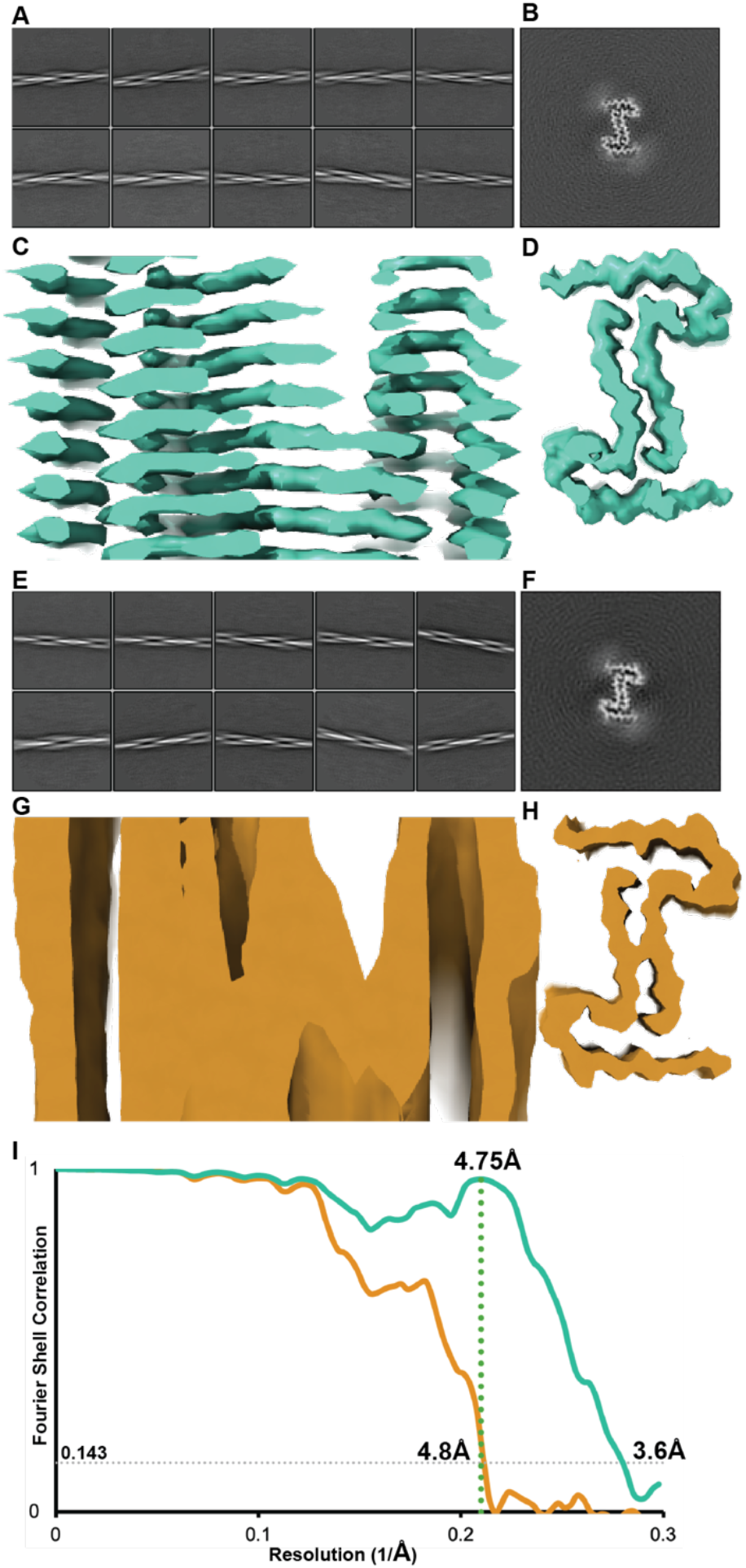
Filaments without strong FOM signals do not yield high-resolution reconstructions. Filaments that were selected by the auto-picker based on strong 4.75 Å FOM signals give good 2D class averages **(A)** and a 3D reconstruction **(B-D)** to resolutions beyond 4.75 Å (**I**). Filaments that were traced manually in the micrographs because they did not have strong FOM signals also yield good 2D class averages **(E)**, but their 3D reconstruction does not extend beyond 4.75 Å (no FOM in ochre, FOM in teal) **(F-I)**.

### Amyloid-specific Blush regularisation prevents overfitting in difficult refinements

To illustrate the advantages of the amyloid-specific Blush regularisation, we compared refinements without Blush, with standard Blush regularisation ^40^ and with amyloid-specific Blush regularisation for three datasets: the PAD12 tau PHFs in Figure 1, the 2PF^L^ hIAPP filaments in Figure 2 and a data set of recombinant PAD12 tau filaments that were assembled *in vitro* using tau seeds from the brain of an individual with familial Pick disease ^43^. For each data set, the three refinements were started from the same 3D initial model as generated by the relion_helix_inimodel2d program.

Because tau PHFs have relatively large, ordered cores, their cryo-EM images are typically of relatively high signal-to-noise ratios, especially when filaments have been assembled from purified protein *in vitro*. Therefore, refinements of recombinant PAD12 tau PHFs suffer little from overfitting, and the maps obtained from all three types of refinements are of similar quality (Figure 5A).

**Figure 5:**
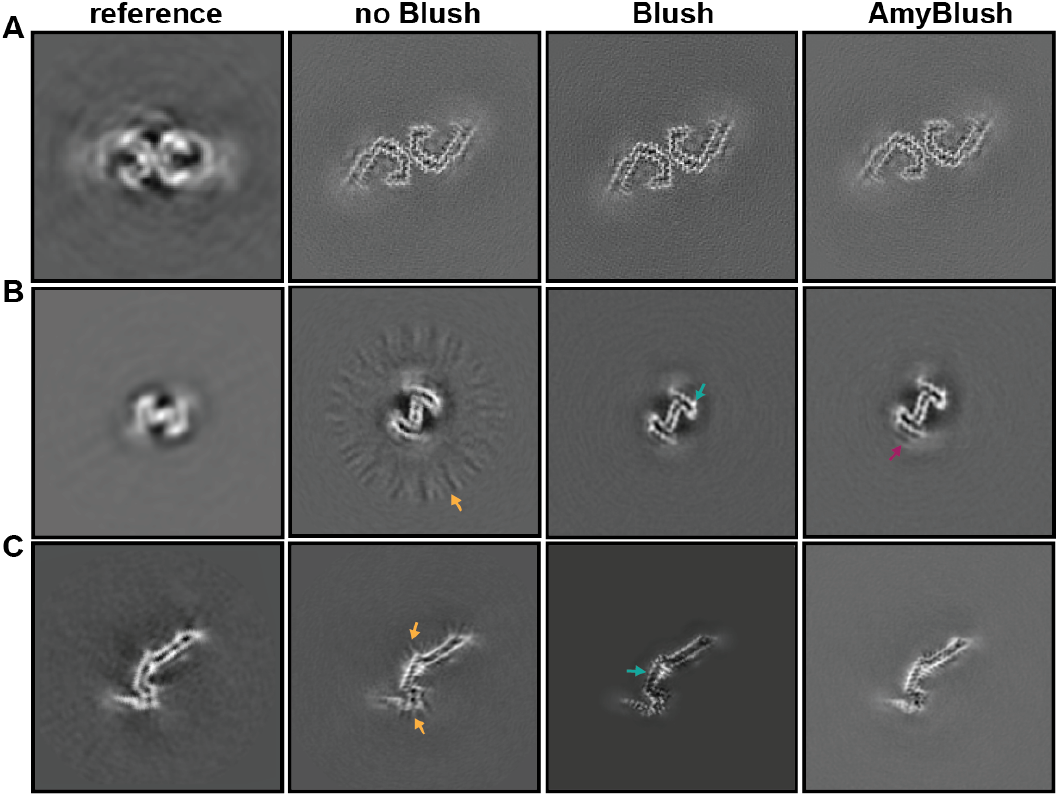
Amyloid-specific Blush regularisation improves difficult refinements. From left to right: slices through the initial model, a map refined without Blush, with standard Blush and with the amyloid-specific Blush for **(A)** the PAD12 PHF dataset; **(B)** the hIAPP dataset; **(C)** the data set on D252V PAD12 tau seeded with filaments from the brain of an individual with familial Pick’s disease. Teal arrows show dotty appearance of the main chain, orange arrows show streaks in the solvent that are indicative of overfitting, and the magenta arrow shows additional features observed for different IAPP filament types.

For the hIAPP filaments, which have a smaller ordered core, refinement without Blush regularisation does suffer from overfitting, as indicated by the radial streaks of density in the solvent region (orange arrow in Figure 5B). For this data set, both types of Blush regularisation remove most of the over-fitting, although the map obtained with amyloid-specific Blush regularisation more clearly showed the presence of additional types of filaments in the data set than standard Blush regularisation (see weak density indicated with a magenta arrow in Figure 5B).

The data set of the Pick-seeded PAD12 tau filaments is the most difficult of the three, possibly because its relatively long and thin ordered core gives rise to wide projections with low signal-to-noise ratios. Despite the use of a tight solvent mask, refinement without Blush regularisation again shows radial streaks in the remaining solvent region, indicating it suffered from overfitting. Refinement with standard Blush regularisation removes the streaks in the solvent region, but the resulting density for the ordered core appears as disconnected dots along the main chain (teal arrow in Figure 5C), and there is density for only a few side chains. Refinement with amyloid-specific Blush regularisation yields a map with connected main-chain density and better side chain densities (Figure 5C).

## Discussion

We present new image processing tools for cryo-EM structure determination of amyloid filaments in RELION-5.1: scripts for fully automated micrograph preprocessing, including a new filament picker; a bi-hierarchical clustering algorithm for classifying distinct filament types; and an amyloid-specific denoiser for Blush regularisation. While the filament clustering approach was already part of RELION-5.0, it had not been formally described yet. We demonstrate the usefulness of these tools on various experimental data sets.

The automated filament picker exploits the 4.75 Å repeat signal that is characteristic of well-ordered amyloid filaments. Thereby, the picker becomes insensitive to filaments that are not amyloids, such as actin filaments, which are sometimes mistaken for amyloid filaments in micrographs on brain or cell-derived samples. Its implementation inside RELION Schemes allows fully automated pre-processing of cryo-EM micrographs, as illustrated for a relatively straightforward data set of PAD12 tau PHFs and a more challenging data set of hIAPP S20G filaments. In both cases, 2D class averages were obtained without user supervision, and the corresponding particles yielded high-quality reconstructions. Although not used in these examples, the Schemes allow on-the-fly processing by iterating through the data while they are being acquired. We envision that standardised pre-processing procedures will increase objectivity and aid high-throughput cryo-EM structure determination efforts, e.g. to find assembly conditions in model systems that replicate the structures from human tissue ^44^.

To our surprise, some of the filaments in the hIAPP data set did not show strong signals at 4.75 Å in their Fourier transforms. Consequently, the auto-picker did not pick these filaments. The observations that these filaments, when picked manually, gave rise to recognisable 2D class averages, but 3D refinements got stuck at 4.8 Å suggest that these filaments are not as well ordered as those that are picked automatically. Therefore, for the purpose of high-resolution reconstruction, the new picker outperforms manual picking. It remains unclear why these filaments are not well ordered. It could be that they became damaged during grid preparation, or that less ordered filaments are an intermediate state in the assembly of these filaments *in vitro*. The latter possibility could be investigated further by a time-resolved analysis of the repeat signals of filaments in the micrographs.

Our approach for selecting different filament types based on 2D classification is similar to the CHEP algorithm ^24,25^, but uses bi-hierarchical clustering on the filaments and the 2D class assignments instead of k-means clustering and PCA. Both approaches use the observation that the structure of the ordered core likely does not change much within individual filaments. The 2D classification of individual particle images is prone to stochasticity due to the high levels of experimental noise, and segments of different filament types may look similar resulting in mixing of filament types in individual 2D classes. Classification of entire filaments using 2D class assignments is more robust than at the individual particle level, as the totality of 2D class assignments along a filament serves as an aggregate signal. Moreover, the bi-hierarchical clustering allows the generation of intuitive displays, in which distinct filament types with varying abundance can be readily identified.

We previously used the new filament clustering tool for the classification of multiple filament types in time-resolved cryo-EM data sets on *in vitro* assembly reactions of recombinant tau ^39^. In this paper, application of the new clustering tool to a time-resolved cryo-EM data set of the *in vitro* assembly of recombinant hIAPP S20G avoided the need for multiple rounds of 3D classification and led to the discovery of two new structures.

When we introduced the original Blush regularisation ^40^, it allowed the reconstruction of the first intermediate amyloid (FIA) in the above-mentioned time-resolved cryo-EM study of recombinant tau ^39^. Comprising only thirty residues, its small, ordered core leads to low signal-to-noise ratios in the images, and hence susceptibility of the refinement to over-fitting. Blush regularisation limited overfitting for these data to the extent that a reliable reconstruction of the FIA could be obtained. However, for other difficult amyloid refinements (not shown), we have observed that standard Blush regularisation may yield reconstructions with “dotty” densities, as exemplified by the tau reconstruction in Figure 5C. It is perhaps not surprising that the standard Blush network does not perform optimally for amyloids. Having been trained only on globular proteins in random orientations, this network may have learnt about general protein features, including the β-sheets that make up amyloid filaments, but it is probably unaware of the repetitive nature of amyloids in the Z-direction. Moreover, it has never seen densities that extend all the way to the top and bottom of the bounding box. By refining the weights of the standard Blush network on 318 amyloid reconstructions and limiting their orientations to those around the helical axis, we aimed to transfer some of the general protein features, while also incorporating new amyloid-specific information into the network. The observation that the “dotty” densities from the original Blush regularisation are no longer present in Figure 5C suggests this objective has been at least partially achieved.

Finally, although the tools described here aim to make amyloid structure determination by cryo-EM reconstruction easier, more objective and more robust, we note that fundamental problems with amyloid refinements getting trapped in local minima remain ^20,26,28^. Therefore, amyloid reconstructions should only be interpreted in terms of *de novo* constructed atomic models if their resolution extends beyond 4 Å *and* if the reconstructed densities show the expected features for protein main and side chains at the claimed resolution, including separation of the main chain densities in the direction of the helical axis. Hopefully, these new tools will be useful in obtaining such structures from more challenging data sets than were previously possible.

## Acknowledgements

We thank Jake Grimmett, Toby Darling and Ivan Clayson for help with high-performance computing; This work was supported by the facility for Electron Microscopy of the Medical Research Council (MRC) Laboratory of Molecular Biology. This work was supported by the MRC, as part of United Kingdom Research and Innovation (UKRI) [MC_UP_A025-1013 to S.H.W.S.]. D.L was funded by a Marshall scholarship; J.S. was funded by a Churchill Fellowship.

## Data and code availability

The cryo-EM maps and models for the 3PF^LU^ and 3PF^LJ^ structures are available from the EMDB (accession numbers 57188 and 57189) and the PDB (29HP and 29HQ), respectively. The source code for RELION-5.1 is publicly available (as branch *ver5*.*1*) from https://github.com/3dem/relion/ and is distributed under the open-source GNU General Public License, version 2.

## Author contributions

All authors performed experiments; S.L. and K.J. supervised parts of the project; overall supervision was provided by S.H.W.S.; all authors contributed to the writing of the manuscript.

## Additional information

For the purpose of open access, the MRC Laboratory of Molecular Biology has applied a CC BY public copyright licence to any Author Accepted Manuscript version arising.

## Supplementary information

**Supplementary Table 1:**
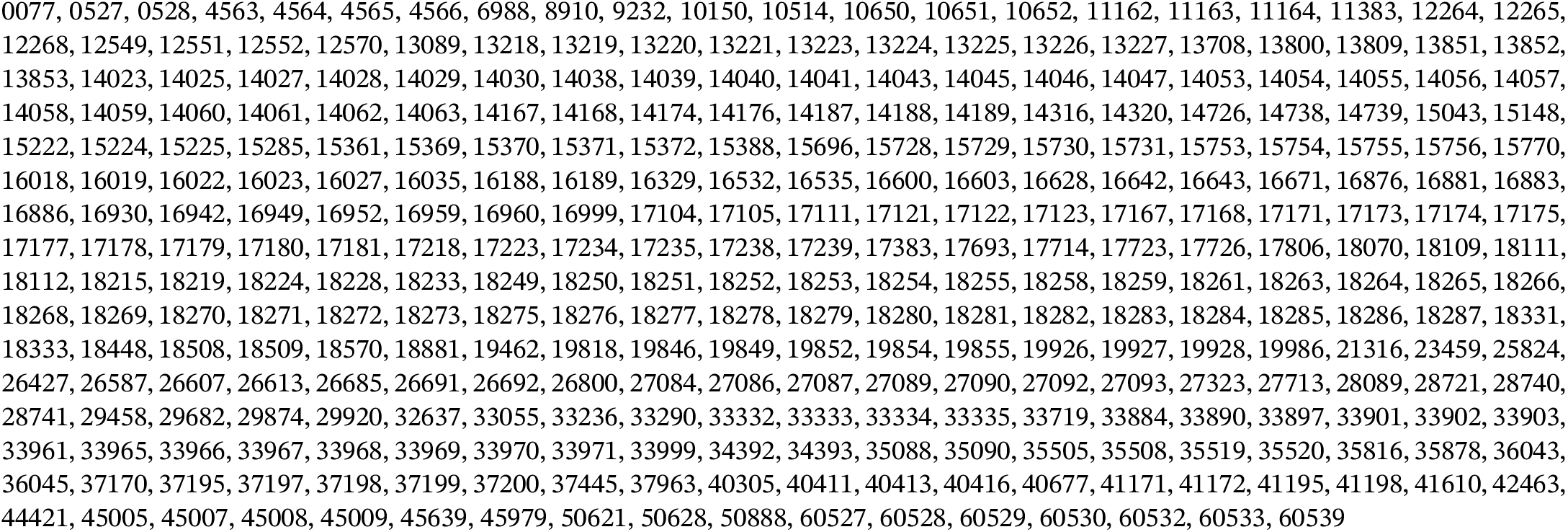
Accession numbers for the 318 EMDB entries that were used for training the amyloid-specific Blush regularisation network:

**Supplementary Table 2:**
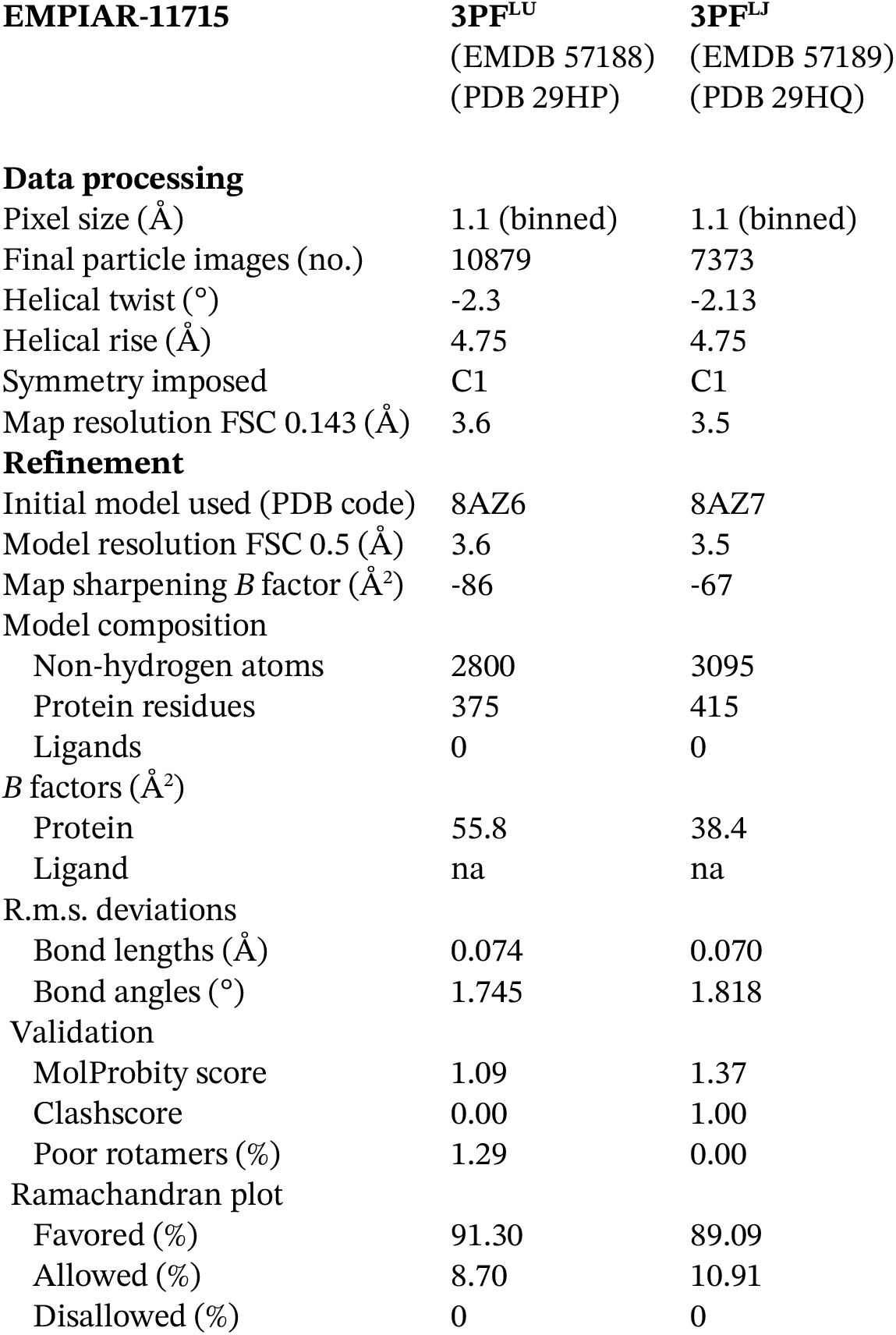
Refinement statistics for 3PF^LU^ and 3PF^LJ^.

| EMPIAR-11715 | 3PF<sup>LU</sup><br>(EMDB 57188)<br>(PDB 29HP) | 3PF<sup>LJ</sup><br>(EMDB 57189)<br>(PDB 29HQ) |
| --- | --- | --- |
| **Data processing** |  |  |
| Pixel size (Å) | 1.1 (binned) | 1.1 (binned) |
| Final particle images (no.) | 10879 | 7373 |
| Helical twist (°) | -2.3 | -2.13 |
| Helical rise (Å) | 4.75 | 4.75 |
| Symmetry imposed | C1 | C1 |
| Map resolution FSC 0.143 (Å) | 3.6 | 3.5 |
| **Refinement** |  |  |
| Initial model used (PDB code) | 8AZ6 | 8AZ7 |
| Model resolution FSC 0.5 (Å) | 3.6 | 3.5 |
| Map sharpening *B* factor (Å<sup>2</sup>) | -86 | -67 |
| Model composition |  |  |
| Non-hydrogen atoms | 2800 | 3095 |
| Protein residues | 375 | 415 |
| Ligands | 0 | 0 |
| *B* factors (Å<sup>2</sup>) |  |  |
| Protein | 55.8 | 38.4 |
| Ligand | na | na |
| R.m.s. deviations |  |  |
| Bond lengths (Å) | 0.074 | 0.070 |
| Bond angles (°) | 1.745 | 1.818 |
| Validation |  |  |
| MolProbity score | 1.09 | 1.37 |
| Clashscore | 0.00 | 1.00 |
| Poor rotamers (%) | 1.29 | 0.00 |
| Ramachandran plot |  |  |
| Favored (%) | 91.30 | 89.09 |
| Allowed (%) | 8.70 | 10.91 |
| Disallowed (%) | 0 | 0 |

